# Test-retest reliability of resting-state EEG intrinsic neural timescales

**DOI:** 10.1101/2024.09.09.611966

**Authors:** Xiaoling Tang, Shan Wang, Xinye Xu, Wenbo Luo, Mingming Zhang

## Abstract

Intrinsic neural timescales (INTs), which reflect the duration of neural information storage within local brain regions and capacity for information integration, are typically measured using autocorrelation windows (ACWs). Extraction of INTs from resting-state brain activity has been extensively applied in psychiatric disease research. Given the potential of INTs as a neural marker for psychiatric disorders, investigating their reliability is crucial. This study, using an open-source database, aimed to evaluate the test-retest reliability of ACW-0 and ACW-50 under both eyes-open and eyes-closed conditions across three sessions. The intraclass correlation coefficients (ICCs) were employed to quantify the reliability of the INTs. Our results showed that INTs exhibited good reliability (ICC > 0.6) at the whole-brain level across different index types and eye states. Spatially, except for the right temporal region in the eyes-open condition, all other regions showed moderate-to-high ICCs. Over 60% of the electrodes demonstrated moderate-to-high INT ICCs under both eyes-open and eyes-closed conditions, with ACW-0 being more stable than ACW-50. This present study comprehensively assessed the reliability of INT under various conditions, providing robust evidence for their stability in neuroscience and psychiatry.

## Introduction

Resting-state electroencephalography (rsEEG) is a non-invasive neurophysiological technique that records the spontaneous neural activity of the brain without specific tasks [1, 2]. It is a critical functional biomarker for evaluating neurological dysfunction and is widely used in psychiatric studies [3,4], including, but not limited to, major depressive disorders [5], schizophrenia [6], addiction [7], and Alzheimer’s disease [8]. Additionally, examining rsEEG oscillatory activity across frequencies may shed light on the function of neural circuits that support a range of cognitive processes, such as attentional control, information processing, and memory [9,10]. Although the brain executes numerous complex cognitive tasks on a daily basis, the neural mechanisms underlying the integration and processing of information across different temporal scales remain poorly understood. Recent studies on neural activity during the resting-state have identified complex, inherent temporal dynamics within the brain’s organization, known as intrinsic neural timescales (INTs) [11–13]. The role of INTs in temporal integration and segregation are crucial for the adaptive processing of external information in the brain, highlighting the importance of temporal dynamics in neural information processing [14].

It is imperative to initially outline the hierarchical structure of INTs across unimodal and transmodal domains to comprehensively understand the functional significance of temporal integration and segregation associated with INTs. Neuroimaging studies employing functional magnetic resonance imaging (fMRI), magnetoencephalography, and related technologies have consistently revealed a pattern in which in unimodal regions, such as the primary visual cortex, exhibit shorter INTs, reflecting their crucial role in rapid and localized neural processing. Conversely, transmodal regions, including the central-executive and default-mode networks, demonstrated longer INTs [11,13,15–17]. This feature is not unique to humans and can also be observed in non-human primates [18,19]. Substantial empirical evidence suggests that this spatial feature may be instrumental in mediating the critical processes of temporal integration and segregation inherent in INTs [20–25]. These findings indicate that, during the processing of sensory stimuli by the brain, information at shorter temporal scales, such as from a note within a melody, is preferentially processed in unimodal regions. Regions with shorter INTs enable separate processing of inputs, enhancing the precision and temporal accuracy of stimulus processing. In contrast, long-lasting stimuli, such as from a complete melody, are processed in transmodal regions, where extended INTs enable the integration of varied sensory information into a cohesive whole. The differential modulation of stimulus information across different timescales highlights the complex mechanisms of the brain involved in optimizing cognitive processing.

Recent studies have demonstrated a strong link between INTs and diverse cognitive functions. Zilio et al. [26] investigated the relationship between brain INTs and sensory information processing. They found that under conditions of sensory impairment, the brain’s INTs were prolonged and shifted toward slower frequencies [26]. Conversely, under conditions of motor impairment, INTs did not change. Further exploration has revealed that in conscious states, INTs negatively correlated with alpha peak frequency, which reflected the temporal precision of sensory input. This correlation was disrupted in an unconscious state [27]. Moreover, INTs have been associated with self-awareness [24,28]; individuals exhibit increased INTs when processing self-relevant information [29].

Recent studies have highlighted the potential of INTs as a biomarker for identifying psychiatric disorders. For example, studies on autism spectrum disorder (ASD) have revealed atypical alterations in INTs [30]. Compared with healthy controls, individuals with ASD showed reduced INTs within the sensory and visual cortices, coupled with increased right caudate INTs. Furthermore, INT values in the sensory and visual cortices were positively correlated with the severity of ASD symptoms severity. Studies in patients with schizophrenia have revealed prolonged INTs during self-relevant tasks compared to healthy controls [31]. Using fMRI, significant reductions in INTs were observed in specific cerebral regions in patients with schizophrenia, including the right occipital fusiform gyrus, left superior occipital gyrus, and right superior occipital gyrus [32,33]. These findings indicate a reduced duration of sensory information storage within the visual cortex and posterior parietal areas. Alterations in INTs have also been observed in individuals with epilepsy [34,35], obsessive-compulsive disorder [36], schizophrenia [32,33,37], Alzheimer’s disease [38,39], major depressive disorder [40], and tobacco use disorder [41].

Recently, there has been a significant increase in research on INTs. Using the ‘Web of Science’ database, a keyword search was conducted with the terms “intrinsic neural timescale*” and “intrinsic timescale*” connected via the Boolean operator “OR”. As depicted in Figure 1, publications on INT research until 2024 demonstrate a steady annual increase, with a notable uptick after 2020, suggesting rising interest in the field.

**Fig. 1.**
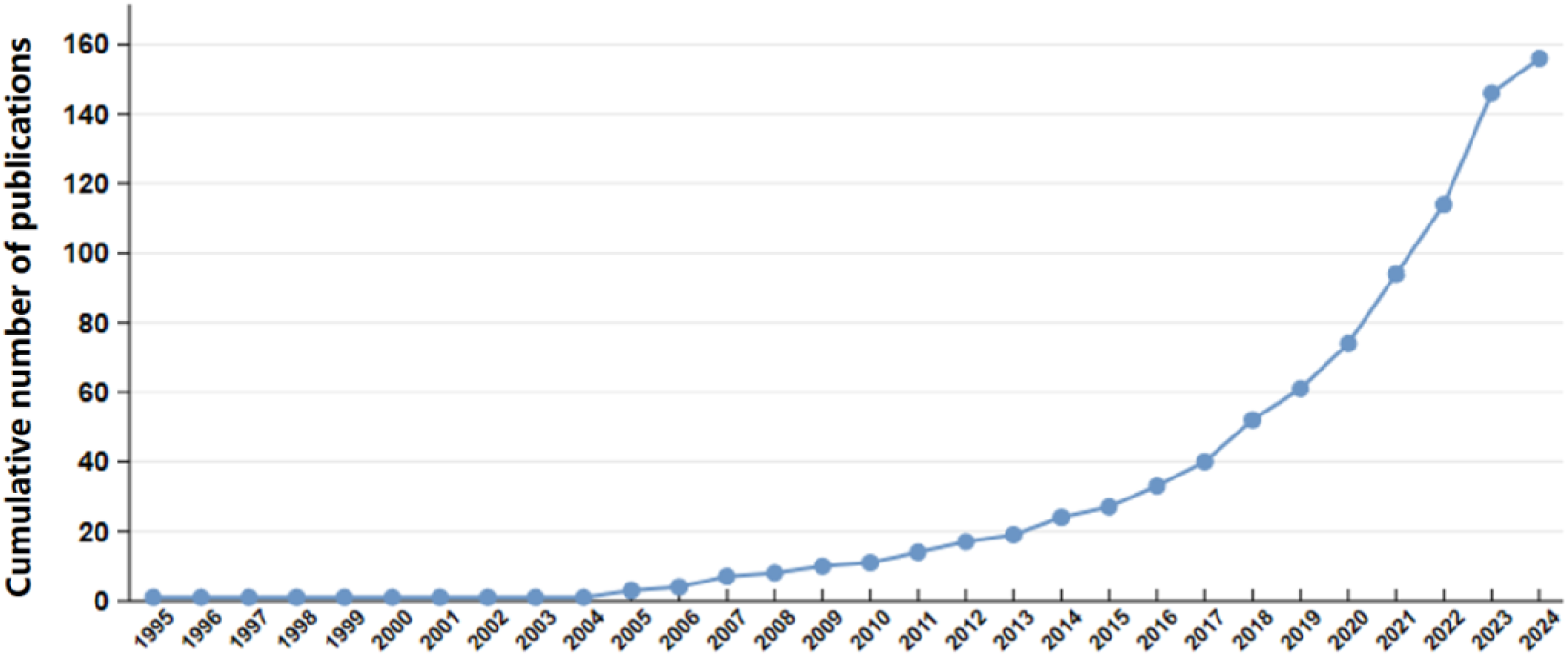
Cumulative number of publications on intrinsic neural timescales.

Despite the widespread application of INT measures in various domains, two major limitations impede their broad adoption and application. First, there has been insufficient research on the stability of INTs. This is particularly crucial when validating INT as a neurophysiological marker because its reliability across repeated measures in the same individual must be thoroughly considered. Low test-retest reliability will reduce the ability to assess the efficacy of treatment interventions [42]. Given this context, this study examined INT reliability. Second, INT measurements are typically reported for autocorrelation window (ACW) lengths, such as ACW-0 [11] or ACW-50 [13,43] (for details, see Methods). However, a detailed analysis of these indices is lacking. Some studies have indicated that ACW-0 encompasses a more extended temporal window and shows a greater predictive capability in differentiating the core and peripheral regions of the brain, compared to ACW-50 [11]. Consequently, this study also examined whether there is a significant difference in stability between ACW-0 and ACW-50, and whether one index is superior to the other.

This study, using an open rsEEG dataset derived from Wang et al. [44], aimed to systematically evaluate the test-retest reliability of the INTs under varying conditions and potential differences between ACW-0 and ACW-50, to facilitate the broader application of INTs in the fields of neuroscience and psychiatry.

## Methods

### Participants and experimental procedures

The data used in this study were obtained from a test-retest resting and cognitive state EEG dataset [44], which is publicly available on OpenNeuro (https://openneuro.org/datasets/ds004148/versions/1.0.1). The dataset includes 60 participants, including 28 males and 32 females (mean age = 20.02 years, SD = 1.88 years). All participants were right-handed; medically screened; had normal or corrected vision; and were free off head trauma, mental disorders, and neurological diseases. Additionally, the participants had not used psychiatric medications within three months prior to the study. On the EEG recording day, the participants avoided alcoholic and caffeinated substances. The participants provided written informed consent following a detailed explanation of the protocol and were compensated for task completion. All experiments were conducted in accordance with the Declaration of Helsinki. The Liaoning Normal University Ethics Committee approved the secondary data analysis.

Eyes-open and eyes-closed rsEEG data were collected over three sessions. The first session was followed by a 90-min interval before the second session, and there was a 30-d interval between the second and third sessions. Participants initially completed 5 min of EEG recording with their eyes open and subsequently completed 5 min of EEG recording with their eyes closed. During the eyes-closed rsEEG recording, the participants were instructed to sit comfortably, close their eyes, avoid engaging in random thoughts, stay awake without falling asleep, and maintain this state for 5 min before opening their eyes. In the eyes-open condition, the participants were instructed to maintain a comfortable position and fix their gaze on a central point on the screen for 5 min, minimizing blinking as much as possible.

### EEG acquisition and pre-processing

EEG data for each measurement were collected using an EEG system (Brain Products GmbH, Steingrabenstr, Germany) with a sampling rate of 500 Hz and 63 Ag/AgCl electrodes. This sampling rate ensured a sufficient temporal resolution to capture rapid neural activity. Electrode placement followed the international 10-20 system, with two electrodes used for recording eye movements and FCz serving as the online reference electrode. Conductive gel was applied to minimize impedance, ensuring all electrode impedances were below 5 kΩ.

Preprocessing of the raw rsEEG data was performed using the EEGLAB toolbox in Matlab2018b. The FCz electrode labels were reinstated to maintain the integrity of the electrode information. The electrode positions were subsequently imported for precise localization. Two electrooculographic electrodes, TP9 and TP10, were removed. The data were subjected to a bandpass filter ranging from 1 to 40 Hz and a notch filter set between at 48 and 52 Hz to eliminate power-line noise. Following established protocols from prior research, the Clean_Rawdata EEGLAB plugin was used to remove flatline channels, low-frequency drifts, noisy channels, and short-time bursts from individual EEG channels, thereby improving the data quality [27,45,46]. Subsequent manual data inspection was performed to identify and exclude artifacts and poor-quality electrodes. Poor-quality electrodes were repaired using the spherical spline interpolation method. Data were then re-referenced to the global average reference and subjected to independent component analysis in EEGLAB to detect and remove artifacts, such as eye movements, blinks, and muscle activity. The data were further segmented into 2 s epochs, and epochs with amplitudes exceeding ± 80 μV were considered as artifact segments and discarded. In the eyes-closed condition, the retention rate of valid epochs was 95.3% (143 ± 6.5 epochs) for the first session, 94.0% (141 ± 7.7 epochs) for the second session, and 93.3% (140 ± 9.2 epochs) for the third session, indicating of high and stable data quality. Likewise, in the eyes-open condition, the retention rates across the three sessions were 94.0% (141 ± 5.6 epochs), 91.3% (137 ± 7.3 epochs), and 92.0% (138 ± 8.6 epochs), respectively, demonstrating comparably high quality and consistency. Participants with an epoch retention below 80% were excluded from further analysis, resulting in a final sample of 58 for eyes closed and 57 for eyes open.

### Calculation of intrinsic neural timescales

The current study employed ACW-0 and ACW-50 to quantify INTs (see Figure 2A). Based on previous relevant studies, a sliding window technique was used to calculate the autocorrelation function (ACF) of the EEG time series with a 20 s window and a 50% overlap [43,47]. The ACF shows the degree of correlation between a signal and its time-lagged counterparts across different time intervals, thus revealing the underlying and intrinsic periodicities and cycles within the dataset [12,14]. Next, the lag when the ACF peak diminished to 50% was determined and defined as ACW-50, whereas the time lag at which the ACF diminished to 0 was denoted as ACW-0 [11,12]. Finally, the mean ACW for all the electrodes was calculated. Open-access Matlab scripts from the “IntrinsicNeuralTimescales” repository on GitHub (https://github.com/Temporo-spatial/IntrinsicNeuralTimescales-) were used for the implementation of the procedures [26].

**Fig. 2.**
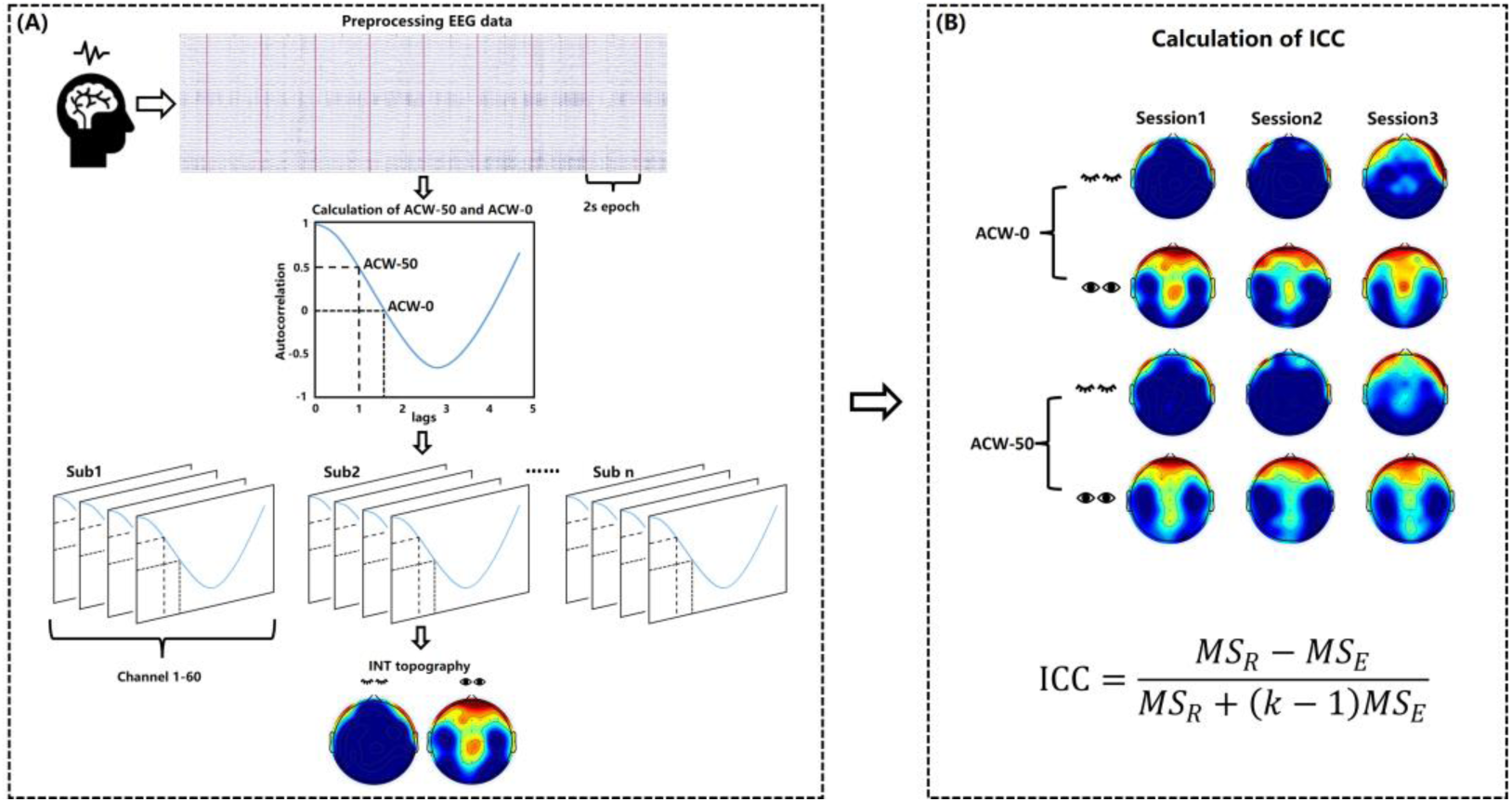
The protocol for data analysis. (A) After preprocessing the raw EEG data and segmenting it into 2-second epochs, ACW-0 and ACW-50 were calculated using autocorrelation function. The whole-brain INT was calculated by averaging INT values across electrodes, and INT topography was plotted. (B) The reliabilities of INT under various conditions were calculated, indexed by the ICCs. ACW: autocorrelation window. INT: intrinsic neural timescale. ICC: intraclass correlation coefficient.

### Calculation of test-retest reliability

The intraclass correlation coefficient (ICC) was used to assess the test-retest reliability of INTs across the three sessions under different conditions. The ICC provides a more precise reliability estimate, reflecting consistency and agreement across measurements [48]. Shrout and Fleiss identified six ICC forms, each stemming from three models and two definitions, demonstrating the complexity and adaptability of the ICC in reliability studies [49]. Consistent with prior EEG reliability research, this study used the ICC (3,1) relation to a two-way mixed-effects model and reported the average of single measurements. The calculation formula is as follow:

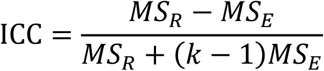

where k (= 3) denotes the number of sessions or measurements. MS_R_ refers to the mean square for rows, which quantifies the variability observed among participants, while MS_E_ represents the mean square of error, reflecting the variability due to measurement errors. The interpretation of ICC values was drawn from existing guidelines in relevant studies. ICC values below 0.4 were considered to reflect poor reliability, those between 0.4 and 0.6 to reflect a moderate level of reliability, those between 0.6 and 0.75 to reflect good reliability, and those above 0.75 to reflect excellent reliability [45,50,51].

The mean INTs of each participant were represented as the average across all electrodes. The whole-brain ICC was calculated based on the average INTs from the three sessions. Additionally, the 60 electrodes were categorized into 11 regions of interest (ROIs, see Figure 4B). ROI 1 comprised electrodes over the left frontal region (Fp1, AF3, F1, FC1, FC3, F3, AF7, F5, FC5, F7); ROI 2 comprised left central region electrodes (C1, C3, C5, CP1, CP3, CP5, P1, P3, P5); ROI 3 comprised electrodes over the left temporal area (FT7, T7, P7, TP7); ROI 4 comprised left occipital electrodes (PO3, PO7, O1); ROI 5 comprised the contralateral counterparts of the right frontal electrodes (Fp2, AF4, F2, AF8, F4, F6, F8, FC2, FC4, FC6); ROI 6 mirrored ROI 2 on the opposite side (C2, C4, C6, CP2, CP4, CP6, P2, P4, P6); ROI 7 was located over the right temporal region (FT8, T8, P8, TP8); ROI 8 corresponded to the contralateral region of the right occipital electrodes (PO4, PO8, O2); ROI 9 comprised midline frontal electrodes (Fpz, Fz, FCz); ROI 10 includes midline parietal electrodes (Cz, CPz, Pz); and ROI 11 comprised midline occipital electrodes (POz, Oz). The ICC of INTs within each ROI was evaluated by calculating the mean INTs of the electrodes within that region. Furthermore, the ICC of INTs at the electrode level was determined by extracting INT data from each electrode and calculating the ICCs across the three sessions. A 95% confidence interval (CI) for the ICCs was also reported.

### Statistics

Paired-sample t-tests were conducted on each electrode to identify those exhibiting statistically significant changes in INTs between the eyes-open and eyes-closed conditions, with a significance level of *p* < 0.05. The Benjamini-Hochberg correction was applied [52] to control the false discovery rate. Furthermore, a two-way repeated-measures analysis of variance (ANOVA) was employed to assess the impact of the index type (ACW-0 and ACW-50) and eye state (eyes-open and eye-closed) on the ICCs of all electrodes. All statistical tests were conducted using SPSS software. Post-hoc comparisons were adjusted using Bonferroni correction to address the issue of multiple comparisons.

## Results

### Topographic distribution of INT

The topographic distributions of the eyes-closed and eyes-open INTs in all three sessions were similar (see Figure 3A and 3B). In the eyes-closed condition, longer INTs occurred in the frontal and temporal channels, whereas other regions showed shorter INTs. In the eyes-open condition, the spatial pattern of INTs was consistent with that reported in previous studies[29,53,54], showing a gradual increase from the unimodal to transmodal areas.

**Fig. 3.**
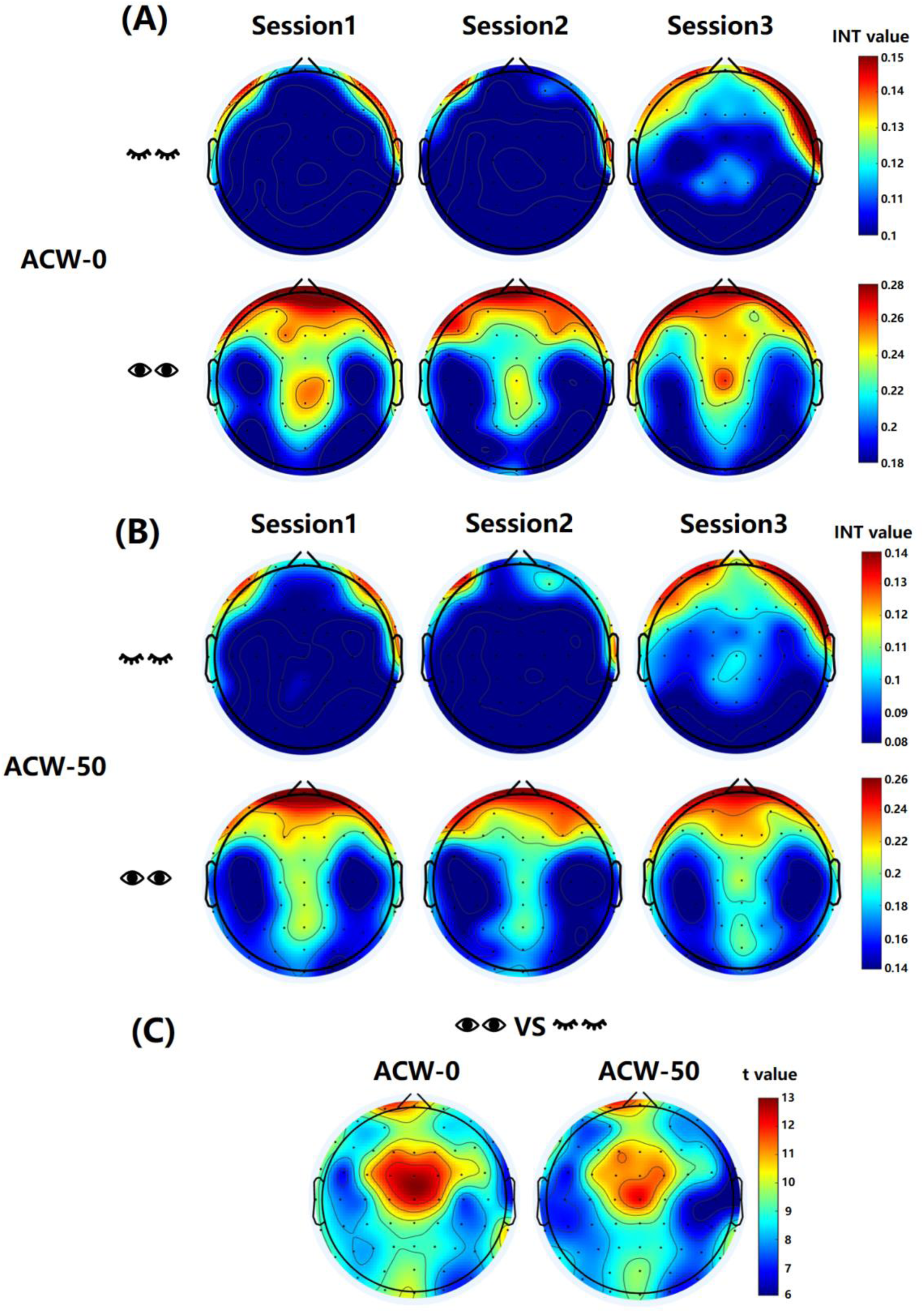
The topographic distributions of intrinsic neural timescales. (A) The topographic distributions of the ACW-0 under all conditions. (B) The topographic distributions of the ACW-50 under all conditions. (C) The topographic distribution difference in ACW-0 and ACW-50 between eyes-open and eyes-closed conditions. ACW: autocorrelation window. INT: intrinsic neural timescale.

Concerning the topographic difference in INTs between the eyes-closed and eyes-open conditions, all t-values of the channels reached statistical significance (*t*_(56)_ ≥ 5.44, *p* < 0.001, after Benjamini-Hochberg corrected), regardless of ACW-0 or ACW-50, especially in the frontal and central regions (see Figure 3C).

### Test-retest reliability of INT Whole-brain ICC

For ACW-0, the ICC values under the eyes-closed and eyes-open conditions were 0.739 (95% CI: 0.632-0.826) and 0.637 (95% CI: 0.503-0.752), respectively (see Figure 4A). For ACW-50, the ICC values under the eyes-closed and eyes-open conditions were 0.685 (95% CI: 0.563-0.787) and 0.632 (95% CI: 0.497-0.748), respectively (see Figure 4A). These results indicate that both ACW-0 and ACW-50 exhibit good test-retest reliability (all ICC > 0.6), regardless of the eye state.

**Fig. 4.**
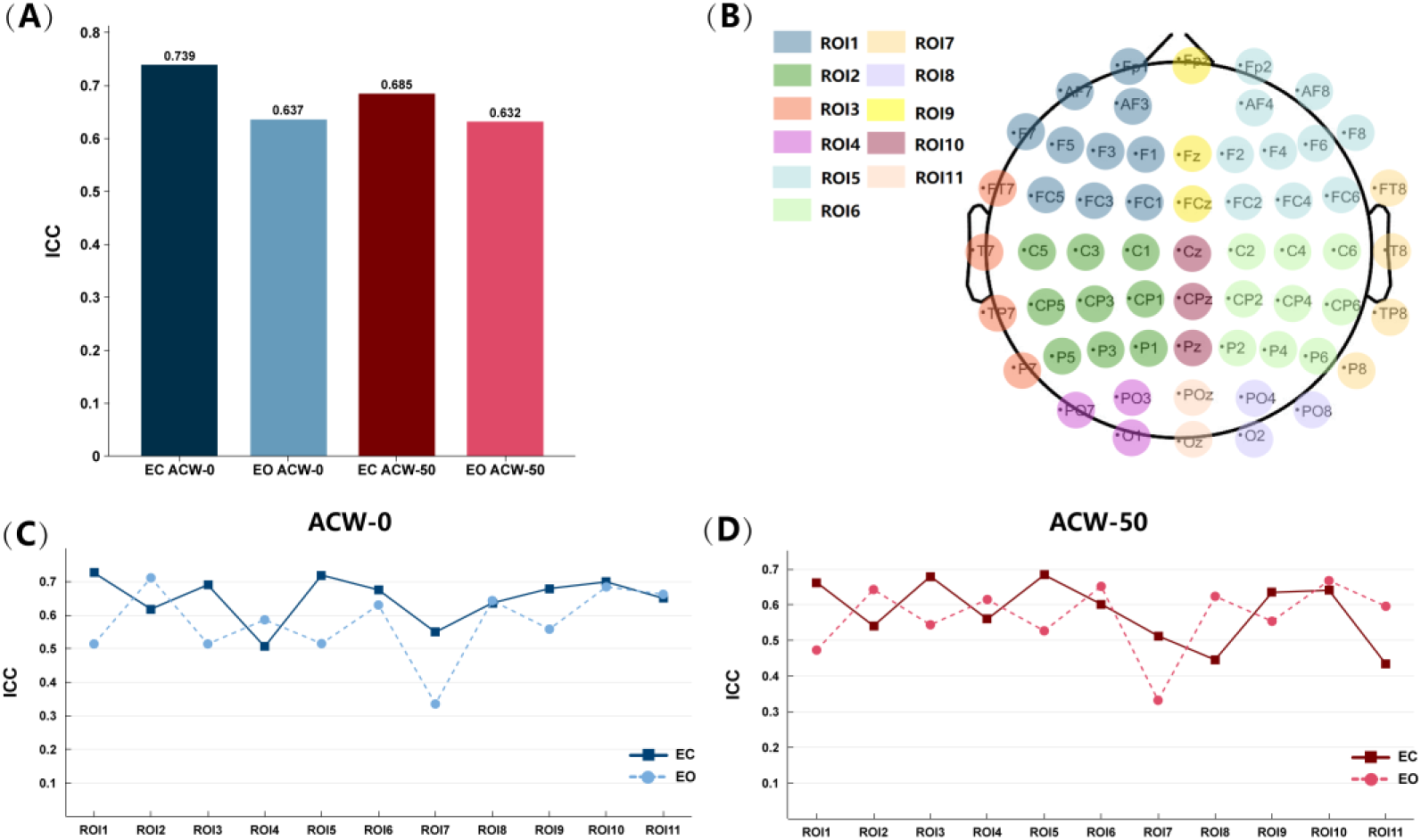
The test-retest reliability results of intrinsic neural timescales. (A) The ICC of whole-brain INT under different conditions. (B) The electrode distribution for eleven regions of interests. (C) The ICC of ACW-0 in different regions of interest and eye states. (D) The ICC of ACW-50 in different regions of interest and eye states. ICC: intraclass correlation coefficient. EC: eyes-closed. EO: eyes-open. ACW: autocorrelation window. ROI: region of interest.

### ROI ICC

For ACW-0, the right temporal ROI 7 showed a relatively low reliability (ICC = 0.335) only in the eyes-open condition, whereas the other regions showed moderate-to-high reliability (0.514 – 0.732; see Figure 4C). In the eyes-closed condition, all regions showed moderate-to high-reliability (0.506 – 0.726; see Figure 4C). Similarly, for ACW-50, the right temporal ROI 7 also showed a relatively low reliability (ICC = 0.332) only in eyes-open condition, while the other regions showed moderate to high reliabilities (0.473 – 0.668; see Figure 4D). In the eyes-closed condition, all regions showed moderate-to-higher reliability (0.434 – 0.684; see Figure 4D). ACW-0 demonstrated higher regional stability than ACW-50. These results indicate that the INTs show acceptable moderate test-retest reliability at both the whole-brain level and in most regions.

### Channel ICC

Under eyes-closed conditions, the ICCs were generally higher and more broadly distributed, particularly for ACW-0 (see Figure 5A). In contrast, under the eyes-open condition, electrodes with higher ICCs were predominantly localized in the central and occipital regions. This pattern was observed for both ACW-0 and ACW-50. Furthermore, the temporal regions, especially on the right side, exhibited comparatively lower ICCs, as did the frontal regions, in the eyes-open condition.

**Fig. 5.**
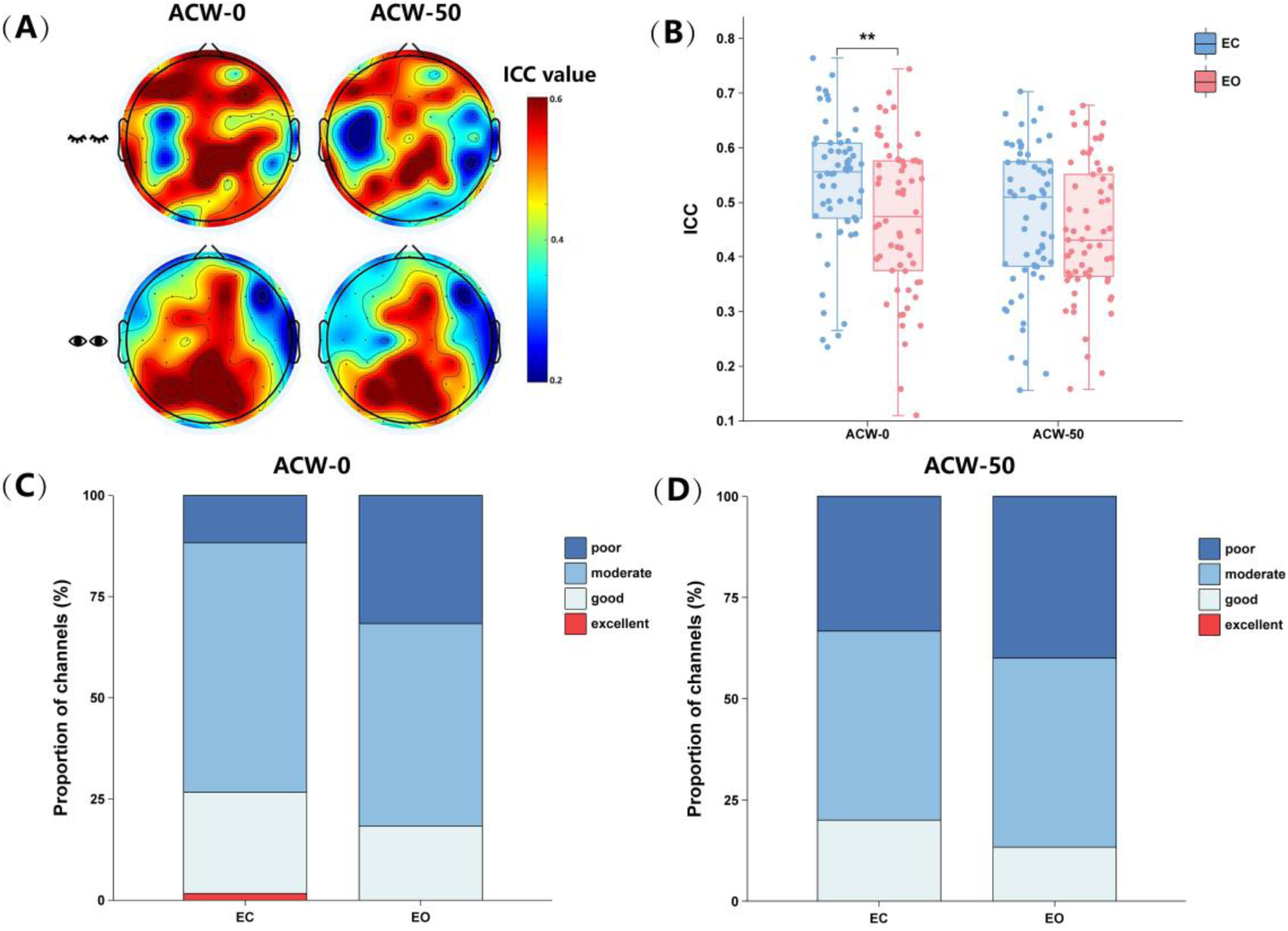
The results of channel reliability for intrinsic neural timescales. (A) The topographic distribution of ICC for intrinsic neural timescales under all conditions. (B) Averaged ICC values for channels under all conditions. (C) The distribution of ACW-0 ICC by reliability ranges. (D) The distribution of ACW-50 ICC by reliability ranges. ICC: intraclass correlation coefficient. EC: eyes-closed. EO: eyes-open. ACW: autocorrelation window. ** *p* < 0.01.

The results of the repeated-measures ANOVA showed significant main effects of index type, *F*_(1,59)_ = 49, *p* < 0.001, η_p_^2^ = 0.454 and eye state, *F*_(1,59)_ = 4.416, *p* = 0.04, η_p_^2^ = 0.07. For the different index types, channel stability was higher for ACW-0 (M = 0.50, SD = 0.13) than for ACW-50 (M = 0.46, SD = 0.13). Additionally, channel stability was greater in the eyes-closed condition (M = 0.50, SD = 0.13) than in the eyes-open condition (M = 0.46, SD = 0.13). More importantly, this result showed a significant interaction between eye state and index type (see Figure 5B), *F*_(1,59)_ = 10.18, *p* = 0.002, η_p_^2^ = 0.147. Further simple-effects analysis showed that the stability of the electrode channels for ACW-0 was greater with eyes-closed (M = 0.53, SD = 0.12) than with the eyes open (M = 0.47, SD = 0.14), *p* = 0.007. However, for ACW-50, there was no significant difference in channel stability between the eyes-closed (M = 0.47, SD = 0.13) and eyes-open condition (M = 0.45, SD = 0.13), *p* = 0.23.

Most channels showed moderate-to-high test-retest reliability (ICCs > 0.4) across the different index types (see Figure 5C and 5D). Specifically, for ACW-0 with the eyes closed, four ICC ranges demonstrated test-retest reliability (poor: 11.7%; moderate: 61.7%; good: 25.0%; excellent: 1.6%). With the eyes open, only three ICC ranges demonstrated the test-retest reliability (poor: 31.7%; moderate: 50.0%; good: 18.3%; excellent: 0%); no channels demonstrated excellent reliability. For ACW-50 with eyes closed, three ICC demonstrated test-retest reliability (poor: 33.3%; moderate: 46.7%; good: 20.0%; excellent: 0%). With the eyes open, three ICC ranges demonstrated test-retest reliability (poor: 40.0%; moderate: 46.7%; good: 13.3%; excellent: 0%). Regarding both ACW-0 and ACW-50, over 60% of the channels achieved moderate-to-high test-retest reliability across the conditions, indicating a notable level of stability.

## Discussion

In this study, we assessed the reliability of INTs using rsEEG data. This study evaluated the test-retest reliability of two distinct INT indices, namely ACW-0 and ACW-50, under two physiological conditions: eyes-closed and eyes-open. Our results showed good reliability of INTs across different conditions. Analysis of INTs at the channels showed that ACW-0 was more stable than ACW-50 under both conditions.

### Topographic distribution of INT

Topographical analysis revealed distinct INT activation patterns under the eyes-closed and eyes-open conditions. In the eyes-closed condition, the INTs at each channel were shorter than those in the eyes-open condition. The topographical distribution of INTs in the eyes-open condition was consistent with previous studies [29,53,54], indicating significant increases in INTs within the frontal, central, and occipital regions. Similar findings have been supported in recent resting-state fMRI studies [55], showing significant INT increases within the visual and auditory cortices during film-viewing tasks. In film viewing, both visual and auditory stimuli require processing, such as lyrics and dialogues. This requires a prolonged temporal window for the integration of sensory information to maintain perceptual coherence and continuity. Moreover, with eyes open, increased neural activity was observed in the prefrontal cortex [56]. The central and prefrontal cortical regions play crucial roles in attention control and other cognitive functions [57,58]. Consequently, an increase in the INTs could enhance brain’s processing of incoming information. This suggests an adaptive mechanism that enhances neural capacity to manage and integrate complex and dynamic sensory inputs essential for cognition. This finding supports the use of INTs to coordinate the temporal integration and segregation of sensory information, highlighting the key role of INTs in neural information processing [14].

### Test-retest reliability of INT

The results indicate that INTs show good retest reliability. Under eyes-closed and eyes-open conditions, the average ICC for ACW-0 was 0.688, and that for ACW-50 was 0.659. In electrophysiological reliability studies, EEG microstates have shown high reliability regarding duration, occurrence, and coverage (short-term ICC = 0.874-0.920; long-term ICC = 0.671-0.852) [59]. In studying rsEEG power spectral reliability [60], studies have reported that the ICCs for the eyes-closed condition regarding relative and absolute power were 0.66 and 0.54, respectively. In the assessment of EEG brain network stability, studies have shown that global graph indices exhibit notable test-retest reliability, with an ICC of 0.59 [61]. In summary, the INT stability outcomes were deemed reliable, indicating a degree of reliability comparable to that of other established EEG measures.

Most ROIs showed adequate stability under both test conditions. Particularly notable was the ACW-0 during the eyes-closed condition, where all examined regions exhibited moderate-to-high retest reliability, with several regions showing an ICC above 0.7. Conversely, stability within the right temporal region during eyes-open condition was lower. Reduced signal stability within the right temporal region may be attributed to interference from electromyographic signals and other forms of noise [62]. The proximity of the temporal region to facial muscles, such as the temporalis and frontalis, coupled with the high activity of the facial muscles, may explain the diminished signal stability [63]. The electromyography noise spectrum, between 20 and 300 Hz, may mask high-frequency brainwave amplitudes, indicating that even minimal noise can significantly affect EEG activity [64]. Research has indicated that the temporal lobe experiences interference from facial and neck muscle activities, even when the eyes are closed [65]. This finding highlights the crucial need to ensure signal integrity in EEG analysis, particularly when examining data from temporal regions.

### Test-retest reliability of INT across channels

The ICC topography across the electrode channels showed that with the eyes closed, the ICC values were higher and more widespread. Conversely, with the eyes open, the ICC values were concentrated in the central and occipital regions, with notably lower values at the frontal and temporal electrodes. The frontal regions are more susceptible to noise artifacts such as blink-induced disturbances during eyes-open conditions [66]. Temporal regions are also susceptible to electromyographic interference [63]. Central electrodes are usually placed near the scalp midline, a location less affected by eye and muscular artifacts. Based on signal-to-noise ratio theorizing [67], it could be hypothesized that the central region’s prominence regarding ICCs may be attributed to its reduced susceptibility to artifacts such as ocular and muscular electrical activities.

Channel stability showed that with the eyes closed, 88.3% of the electrodes exhibited moderate-to-high retest reliability for ACW-0, with an ACW-50 index of 66.67%. In contrast, in the eyes-open condition, electrode stability declined, with the proportion of electrodes exhibiting moderate test-retest reliability for ACW-0 and ACW-50 at 68.33% and 60%, respectively. Both indices showed that over 60% of the channels maintained a moderate level of test-retest reliability across the conditions.

The results concerning the comparative advantages of the two indices revealed a significant main effect regarding the index type. ACW-0 showed greater stability than ACW-50. ACW-50 reflects a shorter time window determined by the duration for the ACF of EEG signal to reduce from peak to half, and may be more susceptible to noise, causing a rapid reduction of the signal to 0.5 [29]. In contrast, ACW-0, which represents a longer time window in which the ACF diminishes from peak to zero, inherently offers better resistance to noise, yielding a higher signal-to-noise ratio. This robustness against noise could explain the greater stability of ACW-0.

Furthermore, this study revealed a significant main effect of eye state. Channel stability was greater in the eyes-closed condition compared than in eyes-open condition, which is consistent with previous findings regarding retest reliability using rsEEG [60,68]. The differential channel stability observed under these two conditions may be attributed to noise interference. The eyes-open state is more susceptible to noise, such as artifacts due to eye movements; for example, blinking rates are elevated when the eyes are open, but in the eyes-closed state, somnolence can lead to ocular rolling [66]. Consequently, the degree of noise influence on the electrode channels varies between conditions, which typically results in greater EEG index stability during eyes-closed than eyes-open states. This variation in noise susceptibility highlights the necessity of considering eye state when interpreting EEG stability. Appropriate noise correction methods should be applied to separate neural signals from noise artifacts to improve the reliability of EEG assessments.

Interestingly, although the eyes-closed condition exhibited greater stability for ACW-0, no difference was observed between the eyes-open and eyes-closed conditions for ACW-50. This finding is consistent with differences between ACW-0 and ACW-50 reported in previous studies. Golesorkhi et al. [11] noted that ACW-0 showed better predictive ability to differentiate between peripheral and core brain regions. Smith et al. [29] observed differential activity across the eyes-open resting state, self-referential and non-self conditions for ACW-0, but not for ACW-50. They posited that ACW-0 had greater variability and provided more information enabling detection of activities across a broader time range [27], and that an increased response to slow-frequency provides distinct detection capabilities in contrast to ACW-50. The reduced sensitivity of the ACW-50 index in capturing subtle differences between the eyes-closed and eyes-open conditions suggests its inferior discriminative capability compared to ACW-0.

The current study also has several limitations. First, the assessment of INT test-retest reliability was confined to scalp-level measurements, excluding source-level analysis. Multichannel EEG-based brain network studies have facilitated the discussion of INT stability from a source-level analysis perspective. Second, the temporal scope of the data was limited; the study spanned only 90-min and 30-d intervals, which limits the interpretation of reliability across longer periods or dynamic changes with increasing intervals. Third, the sample size was small. A larger cohort would enhance the accuracy of the results and facilitate the broader applicability of INT measures.

## Conclusion

In summary, this study used ICCs to evaluate the test-retest reliability of INTs. ACW-0 and ACW-50 exhibited good reliability across all conditions, with particularly high reliability under the eyes-closed condition for ACW-0. ACW-0 showed greater reliability in both the eyes-open and eyes-closed conditions. INTs were found to be a stable neurophysiological marker with adequate test-retest reliability, which highlights the potential of INTs in relation to brain science, cognitive neuroscience, psychiatry, and other fields.

## Acknowledgments

This work was supported by the National Natural Science Foundation of China (32200908 and 32020103008) and the Educational Department Foundation of Liaoning Province (LJKQZ20222360).

## Competing Interests

The authors declare that they have no competing interests.

## Data, Material and Code Availability

The data, material and code are available from the corresponding author upon reasonable request.

## Notes

### Competing Interest Statement

The authors have declared no competing interest.

